# The impacts of almond pollination on honeybee viral dynamics

**DOI:** 10.1101/2025.10.17.678657

**Authors:** Nina A. Sokolov, Graham R. Northrup, Lena Wilfert, Michael Boots

**Affiliations:** Department of Integrative Biology, University of California, Berkeley, Berkeley, CA 94720, USA; Department of Ecology and Evolution, University of Chicago, Chicago, IL 60637, USA; Institute of Zoology, University of Regensburg, 93053 Regensburg, Germany

**Keywords:** Virome, honeybee, virus, almond, pollination

## Abstract

Seasonal aggregation of hosts can rapidly reshape microbial and viral communities, with consequences for disease dynamics and spillover risk. Each year, millions of honey bee (*Apis mellifera*) colonies experience a mass human-mediated intercontinental ‘migration’ to California’s Central Valley to pollinate most of the world’s almond supply. Clearly, this ‘mass mixing event’ with hives from across the country has the risk of spreading highly virulent pathogens, including viruses. It is essential to weigh the benefits of the almond bloom against the risks of disease in honeybees, which may also affect native pollinators. We conducted an observational longitudinal RNA-seq study of colonies from a commercial beekeeping operation before, during, and after almond pollination, compared with non-migrating control colonies. We found that viral diversity increased in honeybee colonies during and directly after the bloom; however, it returned to pre-bloom levels a month later. The virome community composition also became more uniform between hives after the bloom. Hives in closer proximity had more similar viromes. This spatial variation suggests that inter-colony drift is a potential transmission route. Together, these findings suggest that the bloom increases viral transmission, with no single virus dominating the communities. Instead, a group of viruses (black queen cell virus, Lake Sinai Virus, deformed wing virus) were responsible for community shifts. Although crop bloom increased viral diversity and community homogenization, this effect was short-lived, with viromes reverting to pre-bloom levels once hives left the orchards. These findings indicate that pollination events can transiently restructure viral communities in managed bees.

## Introduction

Migrations are likely to enhance the geographical spread of pathogens, potentially increasing interspecies transmission and spillover risks (Altizer et al., 2011). Each season, the majority of commercially managed honeybees (*Apis mellifera,* Linneaus 1758) in the United States are transported to California’s Central Valley to pollinate the almond orchards (*Prunus dulcis* (Mill.) DA Webb) (Ferrier et al., 2018; USDA-NASS, 2018), described as one of the largest pollination events in the world (vanEngelsdorp and Meixner, 2010). The almonds require animal pollination and begin blooming in February, when few native pollinators have emerged from overwintering diapause (Klein et al., 2012). Therefore, the industry is currently heavily dependent on honeybees, with each acre of almonds requiring two honeybee hives for pollination (Carman, 2011). Honeybees will regularly fly over three kilometers away from their hives to forage (Beekman & Ratnieks, 2000), meaning bees in a large orchard could potentially share forage and interact with hives from across the country. These interactions pose a heightened risk of parasite and pathogen transmission, including the ectoparasitic mite *Varroa destructor* (Anderson & Trueman, 2000) and RNA viruses.

Although there is concern about how migratory practices impact honeybee health, relatively few studies follow the consequences of commercial hive movement for almond pollination on disease emergence (Cavigli et al., 2016; Glenny et al., 2017; Alger et al., 2018; Faurot-Daniels et al., 2020). Nevertheless, this mixing of hives from all over the US runs the risk of spreading the most transmissible pathogen strains throughout the honeybee population. Theory suggests long-distance global transmission events can select for higher levels of pathogen infectivity and virulence (Boots and Sasaki, 1999). Some focal bee viruses appear to be generalists, being found in a wide range of insect species, some of which show active replication (McMahon et al., 2015; Tehel et al., 2016). Though other studies find that only a small fraction of viruses screened have evidence of active replication across multiple species (Doublet et al., 2025). Therefore, we need to assess the consequences of pollination practices on disease, as they may not only harm managed honeybees but also pose a risk of spillover to cohabiting native bees at stopover sites. While it seems likely that this mass migration would lead to considerable mixing, we have little information on how honeybees from different hives interact within mass-flowering crops (MFCs) and on the implications for viral transmission.

Horizontal transmission of viruses can occur indirectly through the environment on contaminated floral resources (Alger et al., 2019; Burnham et al., 2021) or through direct transmission from natal hive members (Chen et al., 2006) and non-natal honeybees through inter-colony drift and robbing (Fries and Camazine, 2001). MFCs can decrease disease incidence by diluting the indirect interactions on flowers due to the sheer abundance of floral resources (Figueroa et al., 2020; Piot et al., 2021). Alternatively, MFCs have also been shown to increase disease incidence, likely via increased contact between susceptible pollinator hosts (Cohen et al., 2021; Piot et al., 2019). This pattern is expected to be pathogen-specific, with previous work showing decreased deformed wing virus (DWV) prevalence associated with increased non-crop bloom, but not for acute bee paralysis virus (ABPV) (Manley et al., 2023).

Drifting is the movement of individuals from one hive to another and can occur after foraging flights due to orientation errors (Neumann et al., 2000). The rate of drifting depends on both the misorientation and the acceptance level of the new hive (Bordier et al., 2017) and can vary based on the density and configuration of the apiary (Bordier et al., 2017; Seeley & Smith, 2015), virus infection (Geffre et al., 2020; Iqbal & Mueller, 2007), or varroa infestation (Kralj & Fuchs, 2006; Kulhanek et al., 2021). Guard bees at the hive entrance only check a small percentage of incoming bees, and acceptance levels have been shown to increase during times of abundant forage (Bordier et al., 2017). The abundance of floral resources in MFCs and the high density of hives in a mass monoculture landscape with limited visual landmarks may increase hive acceptance and bee orientation errors, thereby increasing drift and the potential for inter-hive transmission. Therefore, honeybee viruses are expected to show spatial heterogeneity, with hives closer together sharing a more similar viral community than hives further away in the orchard due to drift (Pfeiffer & Crailsheim, 1998).

Longitudinal studies that monitor individual colonies are rare, with most focusing on apiary-level health rather than on individual hive dynamics and interactions. A handful of studies have looked at honeybee viruses through time in the almonds (Alger et al., 2018; Faurot-Daniels et al., 2020; Glenny et al., 2017), but have focused on specific viruses rather than the entire viral community. To date, fine-scale temporal tracking of viruses throughout the almond bloom has not been conducted, and in general, few studies monitor individual commercial honeybee hive viruses at a transcriptomic scale over time. As such, there is a clear need for enhanced viral monitoring through surveillance testing to understand viral spread between hives and to monitor viral dynamics associated with the mass movement of honeybee hives for crop pollination. This work is particularly timely, as the 2024-2025 season has recorded the highest losses of commercial honeybees in the United States on record, with over 60% losses totaling more than 1.7 million hives. Early work by Lamas et al. (2025) suggests that viruses are contributing to these record-breaking losses, with highly virulent viral strains detected in dead and dying hives.

Understanding the impact of mass pollination events on the virome of managed honeybees is key to the long-term management of pollinator health. In this study, we investigated the impact of almond pollination on the virome composition of honeybees within a single beekeeping operation in California. The temporal changes in the honeybee virome were quantified throughout the almond bloom via a longitudinal survey following honeybee hives across four time points in the Spring of 2021. Sampling was conducted before, during, and after almond bloom in California and was compared with control samples collected from hives that did not participate in pollination. Treatment hives were sampled across six spatially distinct ‘blocks’ to determine the potential impact of honeybee colony drift. A viral time series was generated, and individual hive viromes were examined over time to identify viruses associated with almond pollination. We hypothesized that hives participating in the almond bloom would experience higher viral pressure, leading to increased viral richness and diversity. If contaminated almond pollen were the potential source of viruses for honeybees, this would only be seen at the final time point in the series, as there is a time delay between when pollen is collected, stored as bee bread, fed to larvae, and then for those bees to emerge as foraging adults (Figure 1). We predict that, if direct interaction is the source of these viruses, we would observe the highest values at the second and potentially third time points, during and after the almond bloom, when the largest number of honeybees from different hives would be interacting. We also predict that hives that were closer together spatially would have more similar viral compositions. Tracking individual hives across the almond bloom enables us to capture temporal change in virome composition and test whether this mass pollination event drives viral community homogenization around a few dominant strains or diversification through increased transmission opportunities.

**Figure 1.**
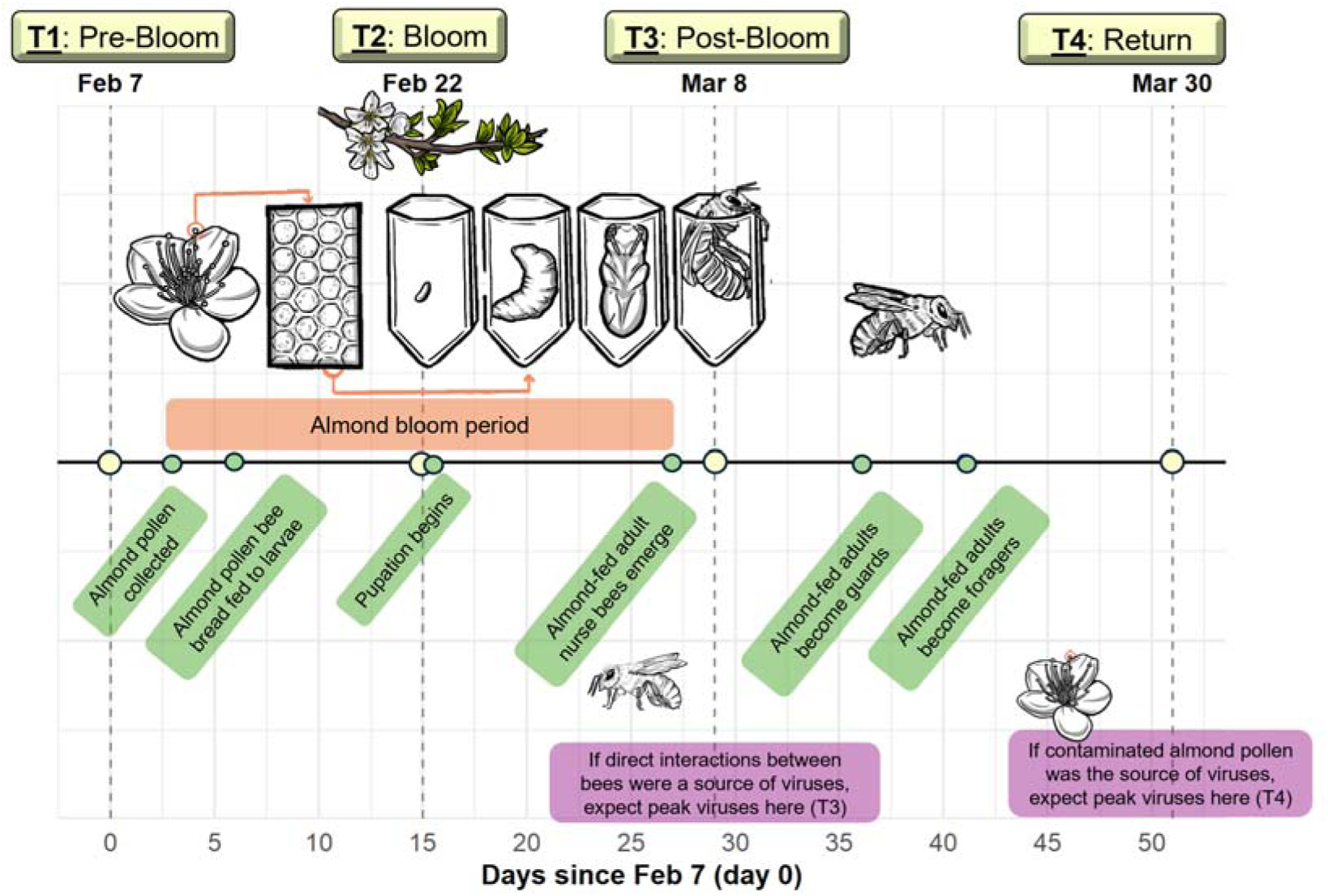
Timeline with days since the first sampling point depicting the almond blooming period and honeybee development. This illustrates the earliest time at which almond pollen would be collected, processed into bee bread, fed to larvae, and then develop into pupae, nurse bees, guard bees, and, finally, forager bees. If contaminated almond pollen were the primary source of viral transmission to honeybees, we would expect the highest viral abundance at the final time point (T4). However, if direct interactions between honeybees were the primary source of viruses, we would expect the highest viral abundance at the time point after all almonds finished blooming (T3).

## Material and Methods

### Sampling and RNA extraction

Sampling was conducted across two almond orchards in California’s Central Valley. Beekeeping operations and management practices act as significant contributors to variation in disease dynamics (Glenny et al., 2017; Martínez-López et al., 2022). To control this variation, we focused sampling on a single California-based beekeeping operation. Honeybee foragers were collected from ‘treatment’ hives that were in an orchard within Arbuckle, CA, and sampled between [02/09/21 – 04/20/21]. A total of 34 treatment hives were sampled during this time, all from the same beekeeping operation. The spatial composition of the orchard consisted of approximately 25 hives per 500 meters in clustered “blocks”. Treatment hives were selected randomly by numbering each hive in the cluster and using a random number generator to select focal hives across six blocks, which were followed for the remainder of the study. All the treatment hives were within a three km radius of one another. Control hives were isolated from direct interactions with almond orchards and sampled in parallel at the beekeeping operation’s home yard in the foothills of the Sierra (Grass Valley, CA), also within a three-km radius of each other (Figure 2a). Due to constraints in the beekeeping operation, the control-hive sample size was small. In total, only three strong hives remained from the almonds and could be used as comparable controls. These hives were part of a breeding management regime to select for *Varroa* resistance. This resulted in samples across four time points: two weeks before the bloom (T1), during the bloom (T2), two weeks after the bloom (T3), and then a final time point when the almond and control hives were brought together three weeks after the bloom (T4) (Figure 2b). Foragers were sampled at the hive entrance using an insect vacuum. Approximately 50 workers were sampled per hive at each time point and placed into 50 ml vials. Samples were frozen on dry ice during transport, with a sub-sample of five foragers per hive placed in cryovials and flash-frozen in liquid nitrogen. Samples were then stored at -80 degrees Celsius on the same day for further processing.

**Figure 2.**
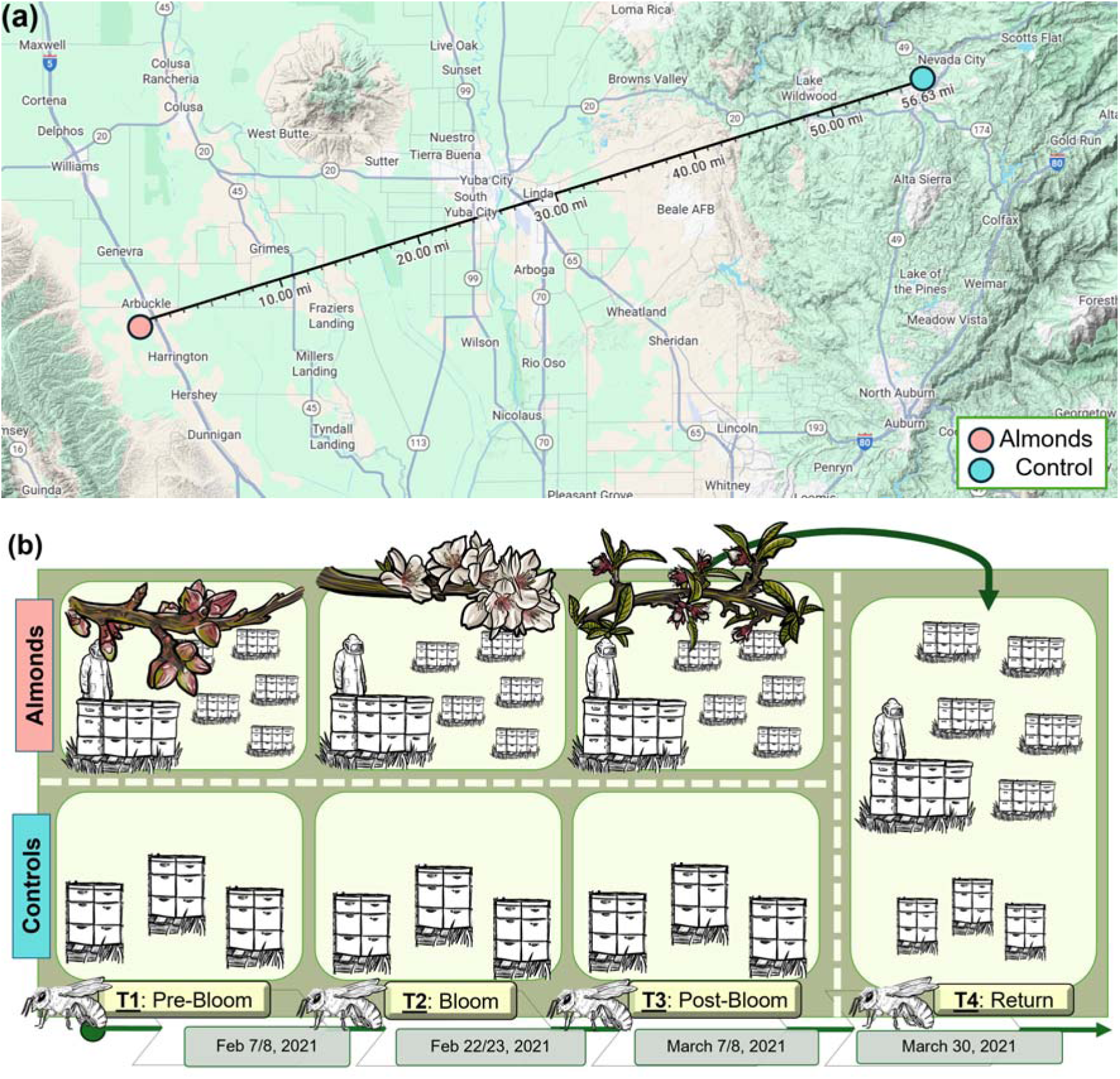
(a) A map of northern California depicting the distance between the almond sites. (blue) and control sites (orange) at approximately a 56-mile distance from one another. **(b) The sampling scheme involved sampling hives in the almonds (top) or controls (bottom) at four time points (February–March 2021).** The first time point was before the almonds bloomed (T1), the second time point was during the almond bloom (T2), and the third time point was after the almonds finished blooming (T3). The final time point involved sampling once the almond and control hives had returned to the home yard.

### Pools for RNA-seq

Five foragers per hive were pooled together by homogenizing tissue in bead-beating tubes containing a single steel ball and zirconia beads in an MPBio FastPrep-24 Classic Bead Mill (speed 4 m/second for 30 seconds; 3 cycles; 20-second dwell) (Runckel et al., 2011). RNA was extracted using standard Trizol protocols (Manley et al., 2020). In brief, RNA was extracted using 1.3 ml TRI Reagent (Invitrogen) and 100 µl of 1-Bromo-3-chloropropane (Sigma-Aldrich), precipitated with 500 µl of isopropanol, washed twice with 1000 µl of 70% EtOH, and eluted in 150 µl of ultra-pure water. DNA was removed from the samples using TURBO DNA-free DNAse Treatment (Thermo-Fisher) and cleaned using an RNA Clean and Concentrator kit (Zymo). RNA concentration and quality were measured using a NanoDrop spectrophotometer (OD260/280 ≥ 2; OD260/230 ≥ 2). RNA integrity of sample pools was tested using a BioAnalyzer Agilent) (s ≥ 4.0, with flat baseline). Samples that passed quality control checks were sequenced on Illumina NovaSeq PE150 platforms at Novogene (Davis, CA). A poly (A) tail enrichment step was performed during library prep to preferentially sequence RNA viruses, which often have long 3’ poly(A) tails, along with host mRNA.

### Bioinformatics pipeline

A total of 131 mRNA-seq libraries were generated from samples that successfully passed quality control. CZ ID was used for the bioinformatic quantification methods (Kalantar et al., 2020). The pipeline included trimming and removal of low-quality reads using ‘fastp’ (Chen et al., 2023) and honeybee host reads using ‘Bowtie2’ (Langmead & Salzberg, 2012) and ‘HISAT2’ (Zhang et al., 2021). After filtration, between 2.5 - 19 million reads remained.

Viruses were classified against the NCBI database using ‘Minimap2’ and ‘DIAMOND’ (Buchfink et al., 2015). Contigs were assembled using ‘SPAdes’ (Bankevich et al., 2012), and reads were mapped against contigs using ‘Bowtie2’. From the alignments, a taxon report was generated that showed taxonomically identified viral taxa of interest, the total number of nucleotide reads (NT r), and reads per million (RPM). Cumulative nucleotide coverage (cumulative_NT) was calculated as:

> Cumulative_NT = Reference length x Coverage breadth x Coverage depth

If contig information was unavailable, the cumulative nucleotide coverage was instead calculated as the number of reads multiplied by the average alignment length (L), although this is an overestimate. Samples were excluded from our analysis if they had < 5 total reads and cumulative NT < 100 (Chen et al., 2023). Low-confidence samples were defined as having >10 reads, cumulative NT > 100, and no contig data, resulting in no available coverage breadth or depth estimates. Moderate confidence samples were defined as read count greater than or equal to ten, breadth of at least 2%, depth greater than 0.1, and cumulative NT greater than 100. High-confidence samples were defined as having at least 50 reads, a coverage breadth of at least 2%, a cumulative NT greater than 250, and a depth greater than 0.5. As viruses were not specifically amplified and our study aimed to capture the full breadth of the viral community, we assessed low, moderate, and high-confidence detections across different analyses, as detailed below. Low-confidence detections were excluded from quantitative analyses. Detections between 10-49 reads were reported but not included in the community-level analysis. A taxonomy table was generated to describe the taxonomic classification of each focal virus. To normalize across differences in sequencing effort amongst samples, we applied a hybrid normalization strategy. If contig-level data were available with coverage depth indicating the average number of reads mapped relative to the viral genome, we calculated normalized depth (avg_norm_depth) as:

> Normalized depth = coverage depth x library size / average library size

For viruses that lacked contig coverage estimates, we normalized with raw read counts as:

> Normalized reads = reads aligned x library size /average library size

From there, both normalized read counts were further converted to relative abundance metrics by dividing the abundance of each virus by the total normalized abundance of all viruses in each sample. To calculate prevalence, we created presence/absence matrices by scoring any non-zero abundance as present and zeroes as absent. For richness and diversity analyses, low-confidence detections with stronger support (≥50 reads and an average alignment length ≥80 bp) were retained, along with moderate- and high-confidence detections. As high- and moderate-confidence detections were supported by contig-level data, both were used to estimate prevalence and quantitative abundance.

### Virus abundance and virome diversity

Analyses were performed in R v4.4.0 (R Core Team, 2014). ‘BioManager’ was used to install ‘Bioconductor’ packages (Morgan & Ramos, 2024). Viral species richness was quantified as the number of different viral species or operational taxonomic units (OTUs) found in each hive at each time point. Richness was modeled using a generalized linear model (GLM) with a Poisson error distribution and log link function. The model included fixed effects of sampling time point, location, and the interaction. Model residuals were examined to confirm assumptions of the Poisson distribution. Additionally, to assess temporal changes in individual viruses within Almond colonies, we fit separate GLMs with binomial error distributions for each virus, using detection (presence/absence) as the response variable and sampling time point as the predictor. Estimated marginal means were calculated using the ‘emmeans’ package (Piaskowksi, 2025), and pairwise contrasts were used to test differences in viral richness between locations and time points, and in viral prevalence between time points in the almonds. P-values were adjusted using the Benjamini-Hochberg procedure to account for multiple comparisons and control for false discovery (Benjamini & Hochberg, 1995).

All plots were visualized using the package ‘ggplot2’ (Wickham, 2016). Absolute abundance was visualized for the viruses with the greatest variation across time points and between locations. Viral diversity within a single sample was assessed from relative abundance data using alpha diversity metrics, including Shannon’s diversity index and Simpson’s diversity index (Magurran and McGill, 2011) and calculated using ‘phyloseq’ (McMurdie and Holmes, 2013). Shannon diversity gives greater weight to rare species, whereas Simpson’s diversity index emphasizes evenness and is therefore more sensitive to common species (Roswell et al., 2021).

We used a linear mixed effects (LMM) model with Shannon/Simpson diversity index as the dependent variable, with fixed effects being “Time” (Levels: T1 = Before bloom; T2 = During bloom, T3 = After bloom; T4 = All back together), “Location” (Levels: Almonds versus Control), their interaction (Time x Location), and “Hive_ID” as a random effect using the ‘lme4’ package. The Shapiro-Wilk Test was used to assess the normality of the distribution of Shannon diversity values. The Levene’s Test for testing homogeneity of variances between groups was performed using the “car” package (Zhou et al., 2023). The model assumptions were quantified using standard tests, including Q-Q, Residual vs. Fitted, Scale-location, and Residuals vs. Leverage plots.

Viral diversity between samples was assessed from each hive’s normalized abundance data using beta diversity metrics, including Bray-Curtis dissimilarity with the ‘phyloseq’ and ‘vegan’ packages (Oksanen et al., 2024). To assess variation in viral community structure across space and time, we conducted a non-metric multidimensional scaling (NMDS) ordination of the Bray–Curtis dissimilarity values. Before ordination, normalized read counts were log-transformed to reduce the influence of highly abundant viruses. Samples with zero total counts were removed. Samples containing only a single detected virus were also excluded, but the data were retained in the overall dataset for other analyses. The resulting ordination was visualized with viral communities color-coded by block, yard, time point, and location to assess clustering. To test for significant differences in community composition, we used Permutational multivariate analysis of variance (PERMANOVA) in the ‘adonis2’ function on the Bray–Curtis distance matrix, with time point, location, and block as predictors (9999 permutations). Data were converted to presence–absence matrices to test effects of community composition using Jaccard dissimilarities computed before applying a similar PERMANOVA model structure.

To assess whether our variables of interest had inherent differences in dispersion values that could be driving differences in beta diversity metrics, we used a permutation test for homogeneity of multivariate dispersions (PERMADISP). The Bray–Curtis dispersion was calculated as the distance of each sample to its hive centroid in multivariate space using ‘vegan’. We performed a SIMilarity PERcentage (SIMPER) analysis to highlight which viruses contributed the most to changes in viral community composition using ‘vegan’. Pairwise dissimilarities were calculated using Bray–Curtis distance, and grouping was based on time point and location. For each pairwise time-point comparison, we extracted the average contribution of each virus to the Bray–Curtis’s dissimilarity. We computed the cumulative contribution to identify the top drivers of compositional change. Viruses contributing to the top 90% of total dissimilarity for each contrast were retained and visualized using bar plots faceted by differences in viral community between time points.

To determine potential co-infection dynamics between viruses, we assessed patterns of viral co-occurrence across hive samples by converting abundance data into a presence/absence matrix. A probabilistic model on the virus matrix data was used to compare observed versus expected co-occurrence frequencies using the ‘cooccur’ package (Griffith et al., 2016) and significant pairs were visualized on a heatmap using ‘pheatmap’ (Kolde, 2018). Finally, normalized log-transformed viral coverage data were analyzed using the packages ‘dplyr’ (Wickham et al., 2018) and ‘viridis’ (Garnier et al., 2024) and plotted using ‘pheatmap’ with hierarchical clustering applied to viral taxa, with grouping by location and time point.

## Results

### Virus prevalence and abundance

We examined the virome composition of 37 hives of honeybees *(Apis mellifera)* from a single commercial operation across four time points in 2021 that either participated in the almond pollination (n=34) or stayed behind at their home yard and acted as controls (n=3). Of those hives, we successfully sequenced and have longitudinal data tracking viral community changes across at least three out of four time points for 29 treatment hives and all three control hives. The remaining eight hive samples did not meet sequencing quality standards for each time point. We identified 28 known insect-associated viruses that met our filters and had more than 2% genome coverage in at least one sample. These viruses mostly belonged to viral families with positive-sense single-stranded RNA genomes (Dicistroviridae, Iflaviridae, and Sinhaliviridae) but also included three Rhabdoviridae with negative-sense single-stranded RNA genomes, one unclassified dsDNA genome, and other unclassified groups. In each hive library, we found between 0 – 8 viruses.

Viral prevalence was assessed for the almond hives and the control hives, as the number of hives found infected with that focal virus over the total number of hives sampled (Figure S1). The viruses with the highest normalized abundance throughout all time points Apis mellifera filamentous virus (AMFV), Apis rhabdovirus, Black queen cell virus (BQCV), Hubei partiti-like virus 34, deformed wing virus (DWV), Lake Sinai virus complex (LSV), sac brood virus (SBV), and uncultured virus 2, which is discussed more below (Figure S2). Some of these viruses in almond hives showed significant changes in viral prevalence over time, suggesting seasonal patterns. We describe the remaining non-significant patterns qualitatively. AMFV prevalence increased during and after the bloom, then dropped to pre-bloom levels at the final time point and was found at relatively high abundance in almond and control hives throughout the study. AMFV prevalence was significantly higher at T2 compared to T4 (p = 0.021) and at T3 compared to T4 (p = 0.036). Acute bee paralysis virus (ABPV) had the highest prevalence before the bloom and was absent in all hives by the final time point. ABPV was detected only in the almond hives and had consistently lower abundance than many of the other viruses of interest.

BQCV was found in both almond and control hives, with prevalence peaking in almond hives after the almonds bloomed, as nearly all hives in the apiary tested positive, but dropped down to pre-bloom levels at the final time point. BQCV prevalence increased from T1 to T2 (p = 0.041) and was significantly higher at T2 compared to T4 (p = 0.034) and at T2 compared to T4 more strongly (p = 0.002). BQCV abundance stayed steady in the almond hives through time and increased at the final time point. In control hives, BQCV was consistently less abundant than in the almond hives. DWV-A and DWV-B were both found at high abundance in the almond hives, with prevalence dropping from the beginning to the end of the study. Both strains’ abundance remained nearly constant before and after the bloom; however, DWV-B abundance dropped sharply at the third time point before rebounding when hives returned to the home yard. DWV-A was identified only as low-confidence reads in control hives at the first two time points and was therefore not included in this analysis.

The most abundant strains of the LSV complex were LSV, LSV-2, and LSV-3, with LSV-2 having the highest prevalence. LSV was higher in almonds than in controls, peaked right after the almond bloom, and returned to pre-bloom levels at the final time point. The control hives had LSV-2 at high prevalence from the beginning, with LSV and LSV-3 absent until the final time point, when prevalence increased to levels like those in the almond hives. LSV-2 was found at a similar high abundance in both almonds and controls pre-bloom but then decreased in the controls at the end of the study. LSV was at a lower abundance in the controls than in the almond hives at the final time point, whereas LSV-3 was at similar levels.

SBV was at relatively low abundance in the almond hives pre-bloom, then increased during bloom, dropped again after bloom, and increased at the final time point when both sets of hives came back together. Prevalence in the almond hives increased consistently through the study. SBV was not found in controls. Uncultured viruses 1 and 2 were found in both almond and control hives at relatively high abundance. Although these viruses have not yet been cultured, previous studies have identified uncultured virus 1 in genomic surveys of varroa (Cornman et al., 2010) and uncultured virus 2 associated with collapsing hives (Cornman et al., 2012). Uncultured virus 1 was high in pre-bloom controls, whereas it was low in almond hives; then, almond hives increased substantially in prevalence after the almonds bloomed, before dropping to pre-bloom levels at the end of the study. Uncultured virus 1 also increased in prevalence from T1 to T2 (p = 0.034). Uncultured virus 2 peaked in abundance in almond hives after the almonds bloomed and peaked in prevalence in the control hives during the bloom.

### Virome diversity

Viral richness was calculated as the total number of unique viral OTUs per hive and time point and modeled using a GLM with fixed effects for time, location, and their interaction. Model diagnostics (residual plots, overdispersion test, and DHARMa simulations) indicated no violations of Poisson assumptions (dispersion p=0.57, zero-inflation p=0.48). Throughout the time series, viral richness increased in almond hives during the bloom (T2) and after the bloom (T3), before returning to pre-bloom levels once the hives left the orchard and returned to control sites (T4) (Figure 3a). We found a significant difference in viral richness between the almond and control samples after the bloom (T3) (almonds > control, p = 0.05). Within the almond location, marginally significant changes in richness were detected between T2 and T4 (p = 0.059), and a significant decrease between T3 and T4 (p = 0.049). The control hives had overall lower viral richness at all time points. Shannon diversity through time was compared between the almond pollinators and control sites (Figure 3b). The distribution of Shannon Diversity values showed a positive skew. A Shapiro-Wilk Test confirmed the data were not normally distributed (p < 0.001). However, when testing model residual diagnostics (Q–Q, residuals vs. fitted, scale–location, and residuals vs. leverage), we saw residuals were approximately normally distributed, with Levene’s test indicating approximate normality and homogeneous variances across groups (F=0.9, p=0.5), supporting the validity of the mixed model (Figure S3). We used the following linear mixed effects model to test the significance of the effect of time, location, and their interaction, along with hive number as a random effect on Shannon Diversity:

**Figure 3.**
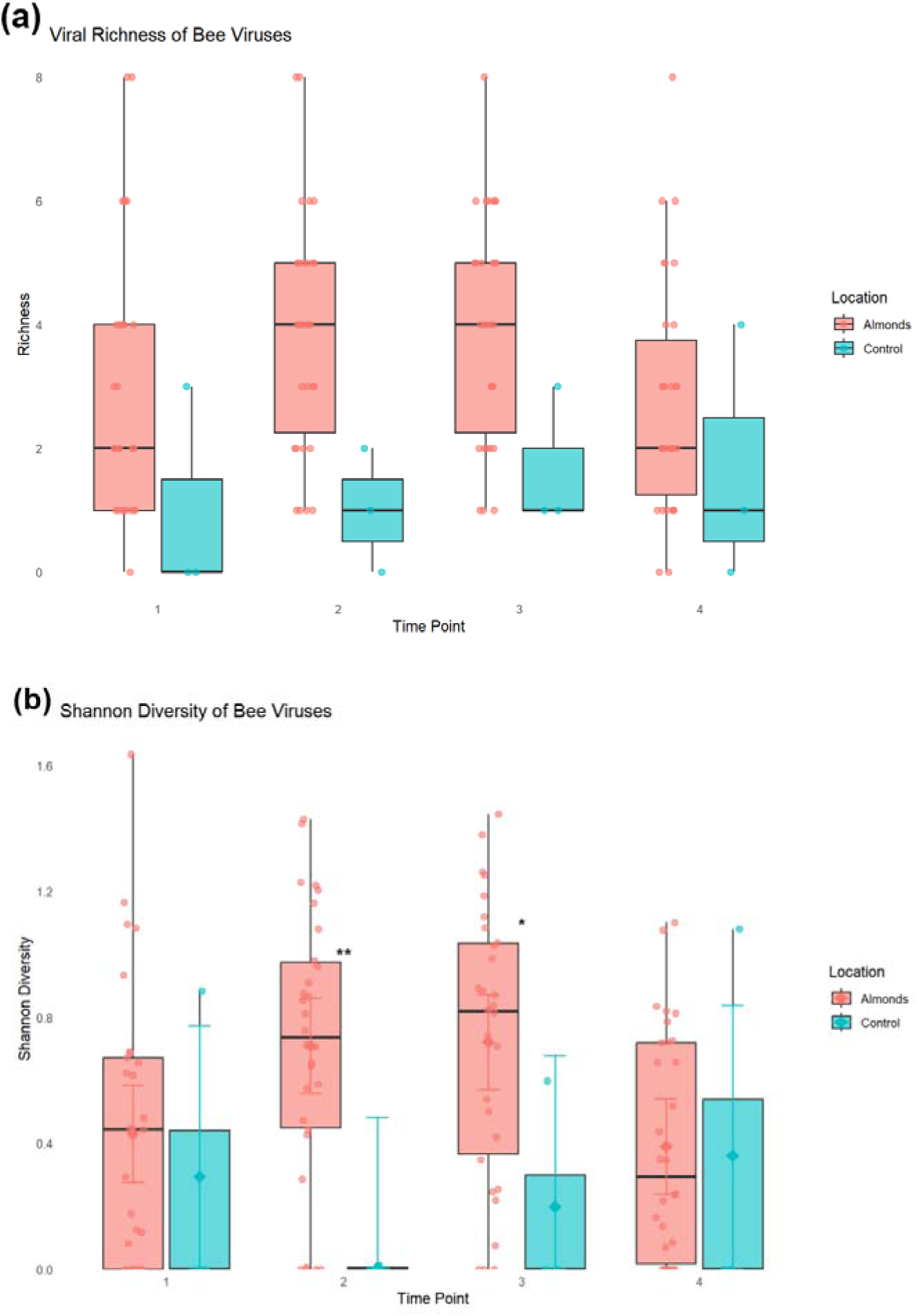
(a) **Viral richness across time in almond pollinating and control honeybee colonies.** Boxplots show the number of distinct viral taxa detected per sample across four time points in almond pollinating hives (pink) and control hives (blue). Individual points indicate viral richness values for each sample. (b) **Shannon diversity of bee viruses across time points and locations.** Boxplots of normalized Shannon diversity indices of viral communities in hives sampled across four time points.

**Model_1** <- m_shannon <- lmer(Shannon ∼ Time_point * Location + (1 | Hive_number), data = diversity_summary)

The model revealed a significant effect of location but not of time on Shannon diversity (Table S1). Residual variance was larger (0.166) than the between-hive variance (0.010), indicating that most variation occurred within hives rather than among them. Post hoc analyses showed that Shannon diversity was significantly higher in almond hives than in controls during mid-season sampling (Table S2). Specifically, diversity increased in almond hives from pre-bloom to the peak of the almond bloom at the second time point (estimate = 0.70 ± 0.25 SE, p = 0.006) and immediately following bloom at the third time point (estimate = 0.52 ± 0.25 SE, p = 0.042). In contrast, early-season (T1) and late-season (T4) samples showed no significant differences between almond and control hives (p >> 0.05). Shannon diversity peaked in almond hives during and immediately after bloom (T2, T3), while controls remained comparatively low, reflecting a transient divergence in viral community structure between locations. When examining changes in viral diversity within hives, we observe variation among hives in whether Shannon diversity increases or decreases during and after the bloom, but most hives return to pre-bloom levels (Figure S4).

Simpson’s diversity index showed similar patterns; however, the control hives at the first time point show low Shannon values but high Simpson values (Figure S5). This indicates that although viral richness is low, no single virus is dominating. This changes at time point 2 in controls, when there are very few viruses present and a single species dominates each sample, resulting in a drop in the Simpson index. In one of the controls, this sample was dominated by ‘uncultured virus 2’, whereas LSV-2 dominated the other sample. Overall, the control hives had variable diversity levels, likely due to the low sample size, but remained low throughout the study. The second time point had the lowest viral diversity, which increased at the final time point when the almond pollinating hives returned.

### Beta Diversity Metrics

To test differences in viral community diversity between samples, we calculated beta-diversity indices and accounted for differences in species abundance using Bray-Curtis dissimilarity. From there, NMDS ordination plots were generated for hives, and a visual inspection revealed no strong clustering of viral communities by block, yard, time point, or location (Figure S6). To better assess patterns of viral abundance across locations and through time, we generated a heatmap of relative abundance for each of the hive’s viruses (Figure 4). We observed clustering of a core group of viruses, including LSV, LSV-2, BQCV, LSV-3, DWV-A/B, and AMFV. We used PERMANOVA to test the interaction among time, location, and spatial block on viral composition, using Bray-Curtis dissimilarity. We found no significant effect of time point, location, or block on community composition (R² = 0.035, p = 0.36) (Table S3). This suggests that when viral abundance information is retained, overall community structure does not differ substantially among groups. However, when we compared presence-absence patterns using the Jaccard-based PERMANOVA, we found significant effects of time (R² = 0.057, p = 0.0003), block (R² = 0.078, p = 0.0027), and location (R² = 0.016, p = 0.0503) (Table S3). This suggests that presence–absence patterns are more sensitive to spatiotemporal differences than relative abundances.

**Figure 4.**
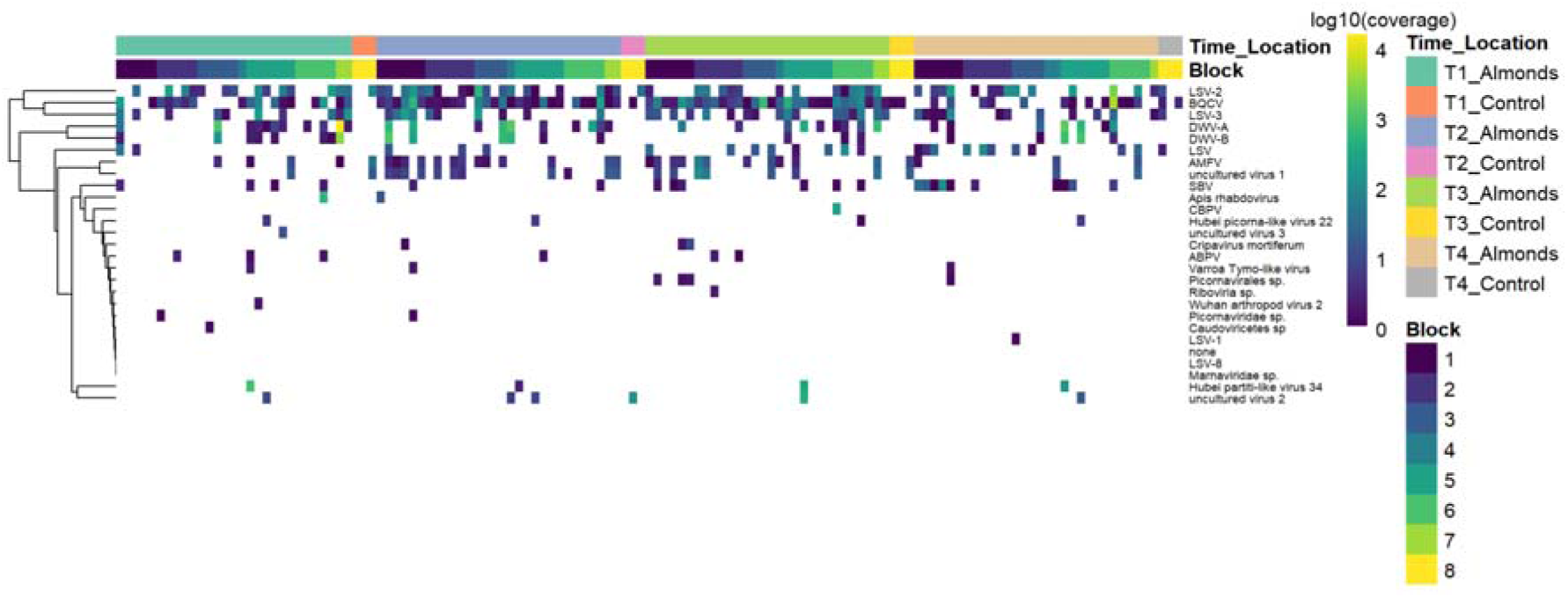
Heatmap of viral community composition across time points, locations, and blocks. Relative abundance values (loglJlJ-transformed) are shown for detected viruses (rows) across all samples (columns). Viruses are clustered by similarity in abundance patterns, while samples are ordered by metadata. Yellow indicates the highest relative log10 abundance, followed by green; darker colors indicate lower abundance.

We calculated beta dispersion within blocks and conducted a PERMANOVA test of homogeneity of multivariate dispersion. We found no significant differences in viral community variation among blocks, time points, or locations (p >> 0.05), indicating that the PERMANOVA assumptions were met. However, when focusing only on almond hives through time, pairwise PERMDISP revealed a significant decrease in dispersion when comparing hives from pre-bloom to directly after the bloom (T1 to T3: F = 4.12, p = 0.037) (Figure 5). This finding suggests that the viromes of almond hives became more similar to one another, or more homogeneous, in their viral community composition after the bloom. However, dispersion returned to levels comparable to those at the first two time points by the end of the study, suggesting that the viral community became more heterogeneous once removed from the orchards.

**Figure 5.**
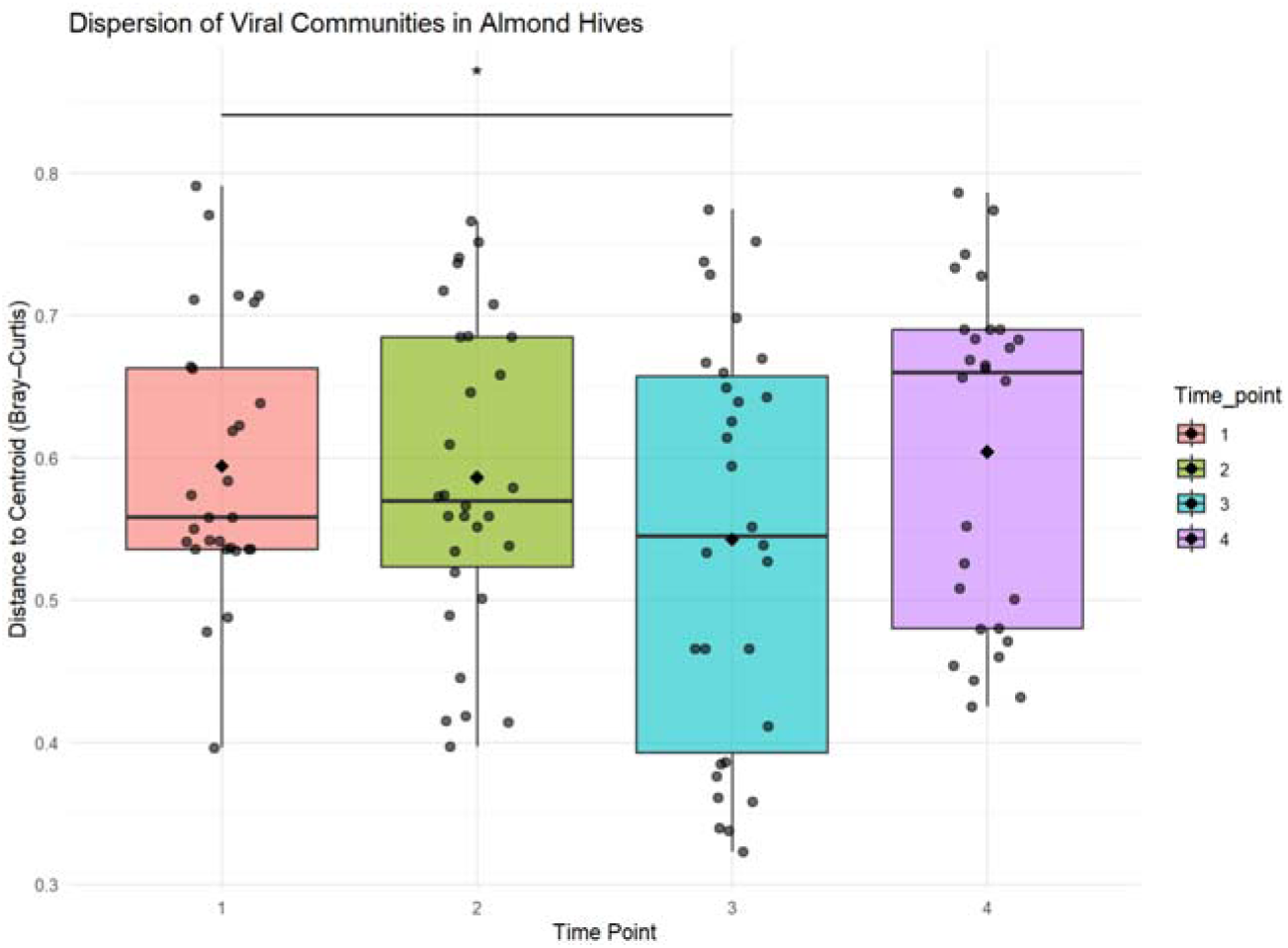
Beta dispersion of viral communities in almond hives across time points. Boxplots of within-group distances to centroids illustrate variation in viral community composition based on Bray-Curtis dissimilarity. Lower values indicate increased community homogenization. Dispersion is significantly different between time points 1 and 3 in the almonds (p_adjusted = 0.03).

SIMPER analysis showed which viruses were the most important contributors to differences in beta diversity between samples. Only viruses contributing to the top 90% of cumulative dissimilarity were shown. A small subset of viruses accounted for most of the dissimilarity between time points (Figure 6). In almond hives, before and during almond bloom (T1-T2), BQCV (18%) and LSV-2 (16%) accounted for the most significant fractions of community turnover. A similar pattern was observed during and after the almonds bloomed (T2-T3), with BQCV contributing the most to changes in the viral community between hives, followed by LSV-2, LSV, and DWV-A. In contrast, control hives showed limited temporal change, with only (T3-T4) yielding significant dissimilarity. BQCV and LSV-2 were again important for this shift in control hives, with uncultured virus 1 and LSV-3 also contributing.

**Figure 6:**
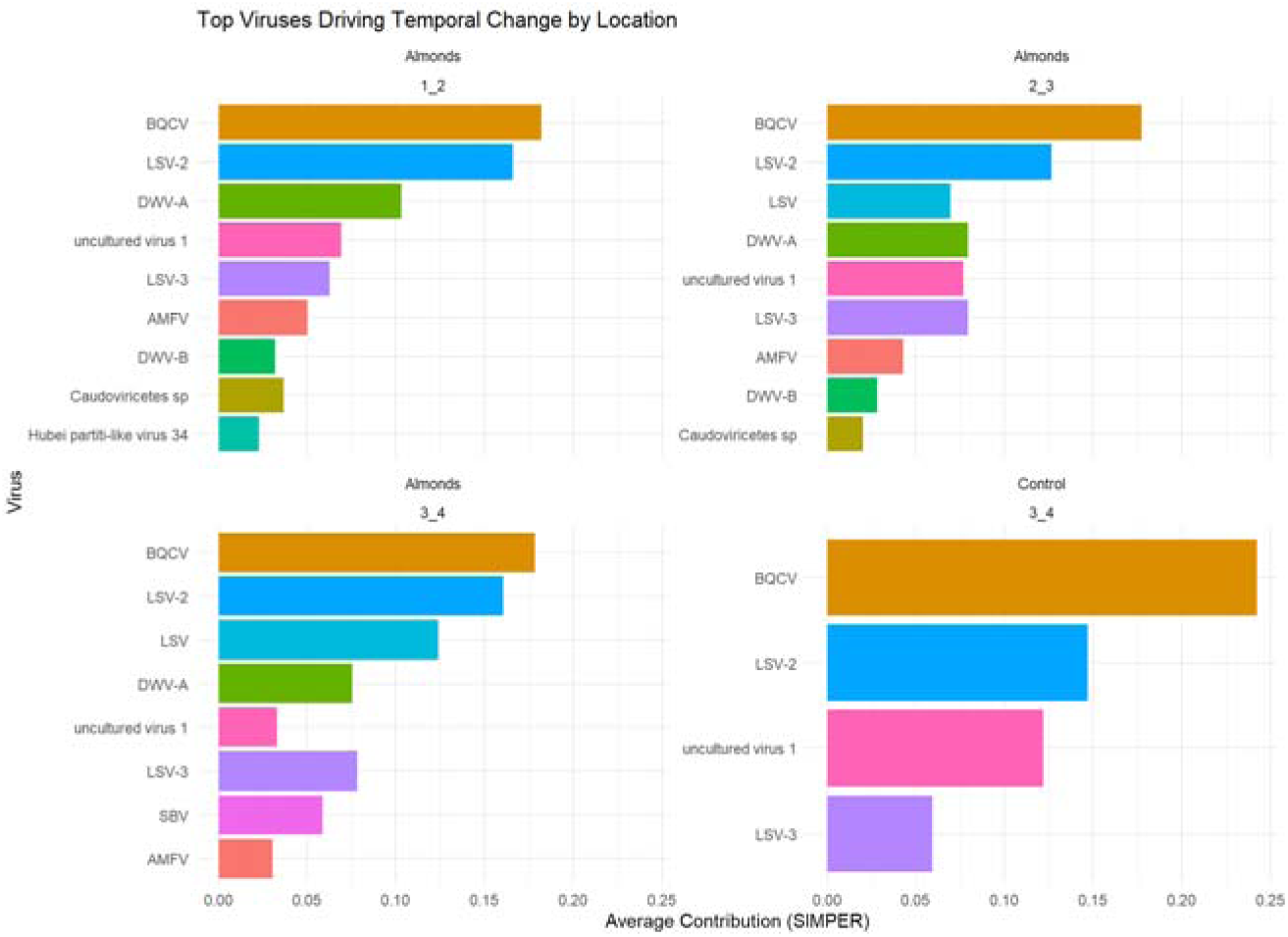
Top viral contributors to dissimilarity in viral community composition between time points. Bar plots show the top viral taxa contributing to differences in beta diversity across all possible time-point contrasts using SIMPER. Each panel shows pairwise comparisons between two time points, with the bars indicating the contribution of each virus to the overall Bray-Curtis dissimilarity. The most influential viruses consistently contributed to community differences across time-point contrasts.

Viral pair co-occurrence patterns were compared using probabilistic modeling to determine how much more or less frequently viruses co-occurred than expected by chance. Significant positive associations were identified after accounting for prevalence and visualized as a heat map for each virus pair (Figure 7). DWV-A and DWV-B showed a positive co-occurrence association (p < 0.001). AMFV strongly co-occurred with uncultured virus 1 (p < 0.001), BQCV, and LSV-3 (p < 0.05). Similarly, LSV-2 exhibited multiple co-occurrence relationships, including with DWV-A, DWV-B, and LSV-3 (p < 0.05–0.001). No viruses showed negative co-occurrence patterns.

**Figure 7.**
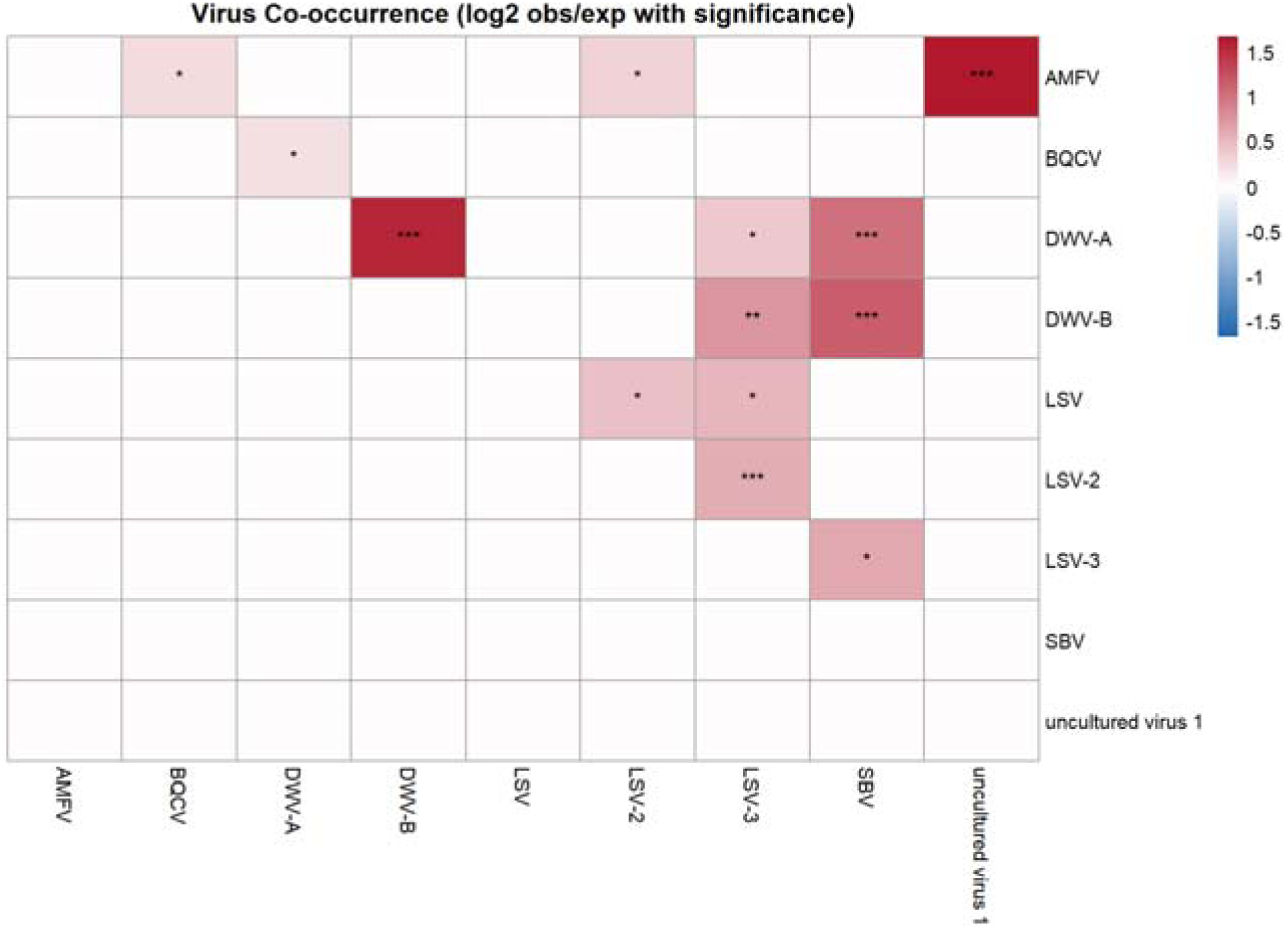
Viral co-occurrence patterns across honeybee hive samples. Heatmap of the loglJ-transformed enrichment of observed versus expected co-occurrence frequencies for virus pairs from probabilistic co-occurrence modeling. Viruses that occurred more often than expected by chance are in red, while co-occurrences that occurred less often than expected by chance were blue. Only viruses that are part of at least one significant pair are plotted. Color intensity is scaled by effect size, and asterisks indicate significance levels (p < 0.05 = *, p < 0.01 = **, p < 0.001 = ***).

## Discussion

Almond orchards can be a high-risk environment for the emergence of virulent pathogens, as they host over half of all commercial honeybees in the US, creating a wealth of susceptible hosts. At the same time, almond pollen is known to be nutritious and thus could bolster honeybee immune function during the bloom as the first natural food source after winter (Loper et al., 1980). The almonds are a unique case in that there is minimal interaction with wild native bees that are not yet active during the peak bloom in February, and pollination is limited by the number of imported hives in the orchards (Reilly et al., 2020). Although honeybees do not directly interact with many wild bee species during the bloom itself (Klein et al., 2012), the bloom’s impact on honeybee viral dynamics could affect native bees when honeybees are moved to their subsequent pollination location. Therefore, it is important to study which viruses are emerging in honeybees because of this pollination event.

We generated an observational time series of viral diversity from a longitudinal RNAseq study of honeybee hives that were shipped to California’s Central Valley to participate in almond pollination in comparison to hives that did not participate in pollination. Our results show a high diversity of viruses, including 28 known insect-associated RNA and DNA viruses. Additionally, evidence of plant viruses and bacteriophages was observed and could be the focus of future studies. Virus prevalence and abundance for focal viruses indicated that almond hives had higher levels than controls, with some evidence of seasonal variation. Our study has shown which viruses increase and decrease in abundance during the bloom and demonstrates significant hive-specific variation in viral dynamics. Viral diversity increased during and after almond blooming at the second and third time points. The highest viral diversity values were observed at the third time point, which we hypothesized was due to direct interactions among bees rather than consumption of contaminated almond pollen (Figure 1). To address this more concretely warrants further investigation, specifically examining the viruses found on almond pollen.

Overall, we observed that viral diversity dropped to pre-bloom levels when almond hives returned to their home yard after completing almond pollination. There was no evidence of specific viruses emerging and dominating the viral community for the almond pollinating bees. Although viral diversity increased during and immediately after the bloom, it returned to pre-bloom levels by the end of the study, suggesting that the effects of participating in almond pollination on bee health may be transient and purge from populations relatively quickly, at least in the studied beekeeping operation. Although viral diversity peaked after the almonds bloomed, viral community dispersion values decreased simultaneously. This means that the almond bloom temporarily homogenizes viral communities, in which either shared floral resources or increased inter-hive contact through drift promotes the spread of a common set of viruses, leading to convergence in community composition across colonies.

We found that virome community composition remained relatively stable across almond and control hives when relative abundances were considered, as evidenced by the absence of significant differences in Bray–Curtis dissimilarity. However, significant effects of time, block, and location were detected when using the Jaccard metric, which uses the presence or absence of viral taxa. The significant temporal effect indicates potential seasonal dynamics. The effect of location also shows a difference in community assemblage between the almond and control hives, which would likely have been even more significant with a larger sample size. The significant block effect indicated that hives located closer together in the orchard had more similar viral communities, suggesting the importance of inter-colony drift. It appears that the presence or absence of rare or transient taxa contributes to detectable shifts in beta diversity over time and space. SIMPER indicated that BQCV and LSV-2 were significant contributors to temporal changes in the viral community in almond hives. In contrast, control hives lacked sufficient viral diversity to yield meaningful contrasts at many time points due to low viral burden. Taken together, these results indicate that while the relative abundance of these dominant viruses may fluctuate over time, the shifts occur within an overall stable virome community. The divergence between these metrics shows that rare taxa drive much of the spatiotemporal turnover in the virome community, even as the dominant viral assemblage remains relatively stable in abundance. Viral infections show several positive associations, with DWV strains and LSV variants frequently co-occurring more than expected by chance, potentially reflecting shared transmission routes. In congregate, these results suggest that honeybee viral communities exhibit subtle temporal and hive-level shifts, yet maintain stable abundance profiles dominated by a few persistent, co-occurring viruses.

BQCV, AMFV, LSV-2, and DWV-A accounted for the greatest change in beta diversity across all time points. LSV is a highly prevalent virus during almond blooms and exhibits high strain diversity, with unknown consequences for honeybees and other bee species (Daughenbaugh et al., 2015). ABPV was detected only in the almond hives, but at lower abundance than many of the other viruses of interest. This virus is of particular concern for potential risks of viral transmission to wild bees, as this is one of the examples of a definitive multi-host virus, as seen in the United Kingdom bee communities (Doublet et al., 2025). Several of the viruses identified in this study have been found in hives across a wide geographic range, but little is known so far about their biology. Hubei partiti-like virus 34 has been found through metagenomic studies in honeybee hives across the world, including the United States (McKeown et al., 2025), Korea (Kwon et al., 2023), and Uzbekistan (Kwon et al., 2024). Uncultured virus 2 and 3 were found in a previous genomic survey of hives experiencing colony collapse disorder, but little is known about them (Cornman et al., 2012). We found individual viruses mostly in the almonds, not the controls. However, this could be explained by low sample sizes. This pattern becomes more pronounced when looking only at high-confidence data. The control hives left behind were the only strong hives that were not used to pollinate because they were part of a breeding regimen focused on varroa-resistance traits. The low disease incidence in controls could then also be a function of their ability to maintain low varroa numbers without chemical treatment (Penn et al., 2022). This was particularly notable for the surprisingly low DWV prevalence and abundance, which are typically high in commercial hives (Molinatto et al., 2025).

Although other studies have characterized honeybee viral prevalence and load before and after almond blooms, it was over a coarser time scale (Alger et al., 2018; Glenny et al., 2017), and only two studies sampled during the pollination event itself (Cavigli et al., 2016; Faurot-Daniels et al., 2020). Runckel et al. (2011) found the dynamics to be sporadic cases of acute infections, in which hives cleared infections after brief periods, with BQCV being the only virus found to persist across time points as a chronic infection. (Alger et al., 2018) also saw BQCV persisting across their experiments. A study that sampled in February and again in March found that hives sampled during the almond bloom had the lowest pathogen prevalence. However, the total number of pathogens sharply increased after the bloom (Cavigli et al., 2016). Certain viruses were prevalent before and throughout the bloom, such as BQCV and LSV (Cavigli et al., 2016). Faurot-Daniels et al. (2020) followed a cohort of 50 individual hives before, during, and after the bloom across the span of a year. They found the highest total number of pathogens per colony a month after almond pollination. This correlated with seasonal buildups in hive populations and increased springtime foraging during the almond bloom and beyond. Although little is known about LSV, they found a correlation between increased LSV2 abundance and weaker colonies, suggesting that LSV is a pathogen of concern. DWV prevalence and load were highest in the fall, LSV was higher in the spring, and BQCV and SBV were both present throughout.

Research has shown that migratory bee colonies experience higher levels of oxidative stress during migration, which in turn may negatively affect the immune function of individual bees (Simone-Finstrom et al., 2016). The existing papers tracking viruses and honeybee health in almonds all followed operations based outside California. The beekeeping operation studied here was in Northern California, approximately 100 km from the almond bloom. The shorter migration distance and potential for local adaptation may mean that this operation experienced fewer stressors than out-of-state beekeepers. The location, along with this operation’s work to breed for varroa resistance in their honeybees, suggests this is a ‘best-case-scenario’ instance for almond pollination. More work is needed with the broader viral community for larger commercial operations that must travel greater distances to reach the almonds and, therefore, face a greater risk of stress.

The critical pollination services that the commercial honeybee hives in the United States provide to the world’s almond crop are commercially essential. The risk of emerging infectious diseases, particularly RNA viruses, is clearly of concern, given the potential for many bee hives across the country to interact. Mass interactions and the spread of viruses across the country seem less likely than we might have predicted. We found that the bloom was associated with a transient increase in viral diversity during and directly after the bloom. Despite bees theoretically being able to interact with hives nationwide during this bloom, viral diversity returned to pre-bloom levels by the end of March. The early mass almond bloom provides the first natural food source for the honeybees emerging from winter, while the pollination services also provide the majority of income for commercial beekeepers. High winter losses before almond bloom are economically consequential for beekeepers but may also differentially cull hives harboring the most virulent pathogens. This suggests that the commercial benefits and early food resources that the bloom provides may outweigh population-level infectious disease risks in this local California-based operation.

## Acknowledgements

We thank Jenn Thompson for helping with sample collection and Sara Herrejon Chavez for helping with RNA extractions. Thank you to Priya Pillai for assisting with RNA preparation. The authors thank the beekeeping operation that collaborated with us during this study, along with the almond growers for providing the study sites. We also thank the bees that were sacrificed for this research. In preparation of this work, the authors used Grammarly to assist with grammar checking and improving language clarity. ChatGPT (version 4) was used to assist with code development, code troubleshooting, and improving wording. After using these tools, the authors reviewed all content and take full responsibility for the conclusions in the manuscript.

## Data Accessibility and Benefit-Sharing

### Data Accessibility Statement

All sequence reads generated in this study will be uploaded onto the NCBI Sequence Read Archive (SRA) database upon publication. Sample metadata will also be available in the SRA submission. The assembled viral genomic sequences will be deposited in GenBank. However, private reviewer access can be provided.

### Benefit-Sharing Statement

The Nagoya Protocol does not apply to this study, as all honeybee samples were collected in the United States. However, benefits were still generated, including sharing these results with the beekeeping community at beekeeping association meetings and conferences.

## Author Contributions

NS and MB designed the project. NS performed the research, collected the samples, conducted the laboratory work, analyzed the data, and wrote the manuscript. GRN assisted with the analysis and statistical testing. LW assisted with analysis and manuscript editing.

## Funding

N.S. was supported by the National Science Foundation (NSF) Graduate Research Fellowship Program. M.B. was supported by the NSF-DEB-2011109 EEID research grant.

## Notes

### Competing Interest Statement

The authors have declared no competing interest.

### Summary of Updates

The manuscript was edited for grammar, clarity, and formatting requirements. No conclusions or results changed between submissions.

