## Supplementary material for "The impacts of almond pollination on honeybee viral dynamics": Supplemntal Tables

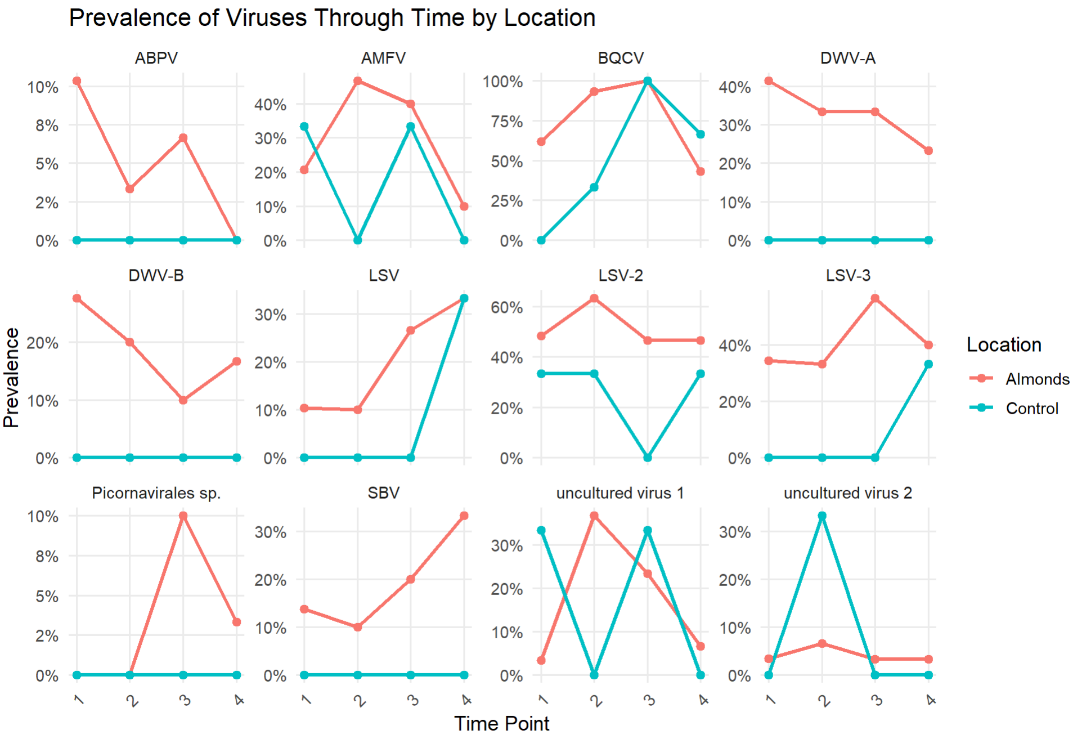

**Figure S1. Viral prevalence over time for honeybee colonies in almond pollination and control groups.** Line plots show, for each time point, the number of hives out of the total found to have each of these viruses in almond (pink) versus control (blue) hives, indicating prevalence as a percentage of positive cases for each virus across the apiary.

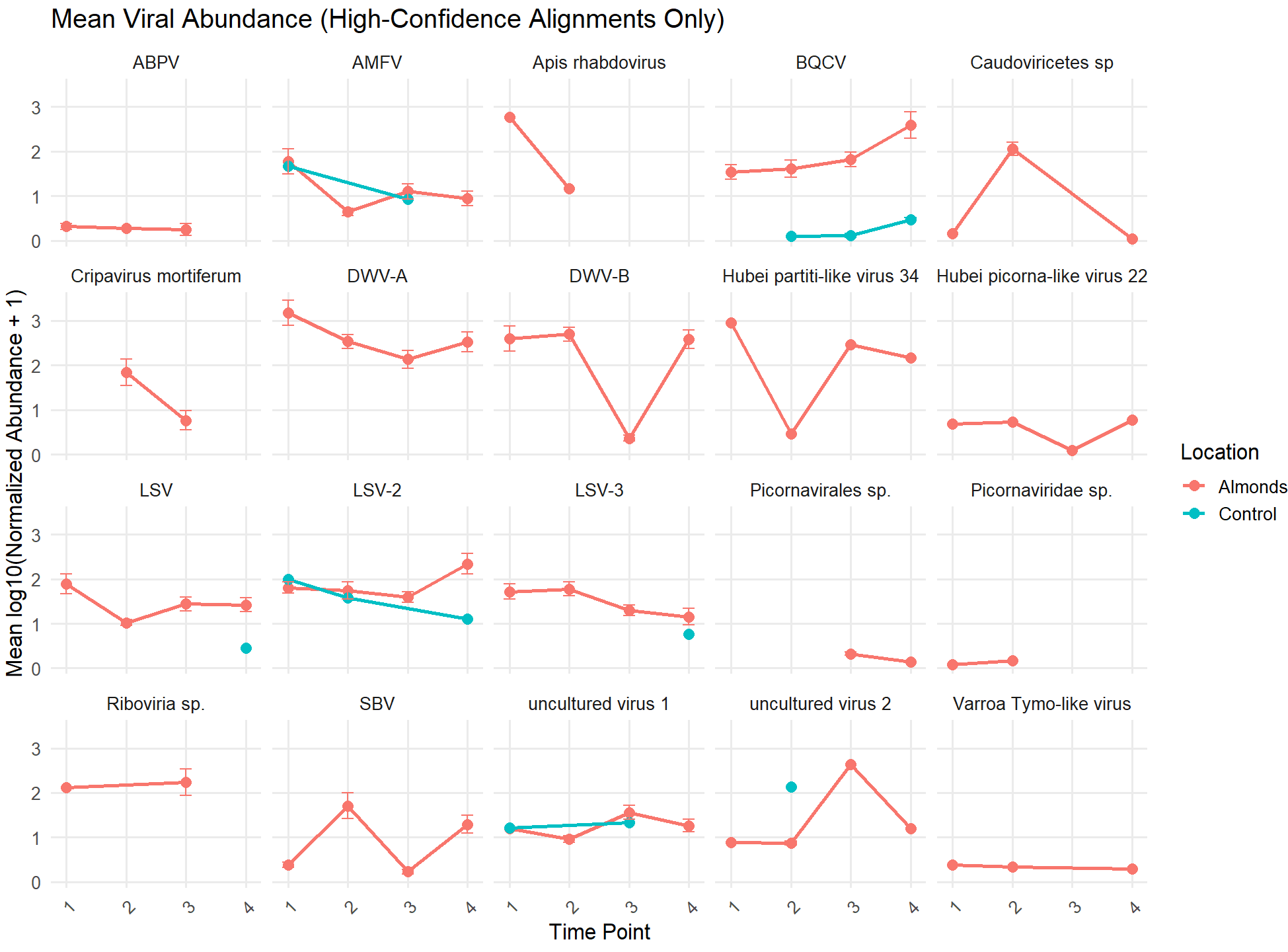

**Figure S2. Mean viral abundance over time for honeybee colonies in almond pollination and control groups.** Line plots show the mean viral abundance (log10-transformed, normalized abundance) for individual viruses over time. Each panel shows the temporal dynamics of mean viral abundance for each group.

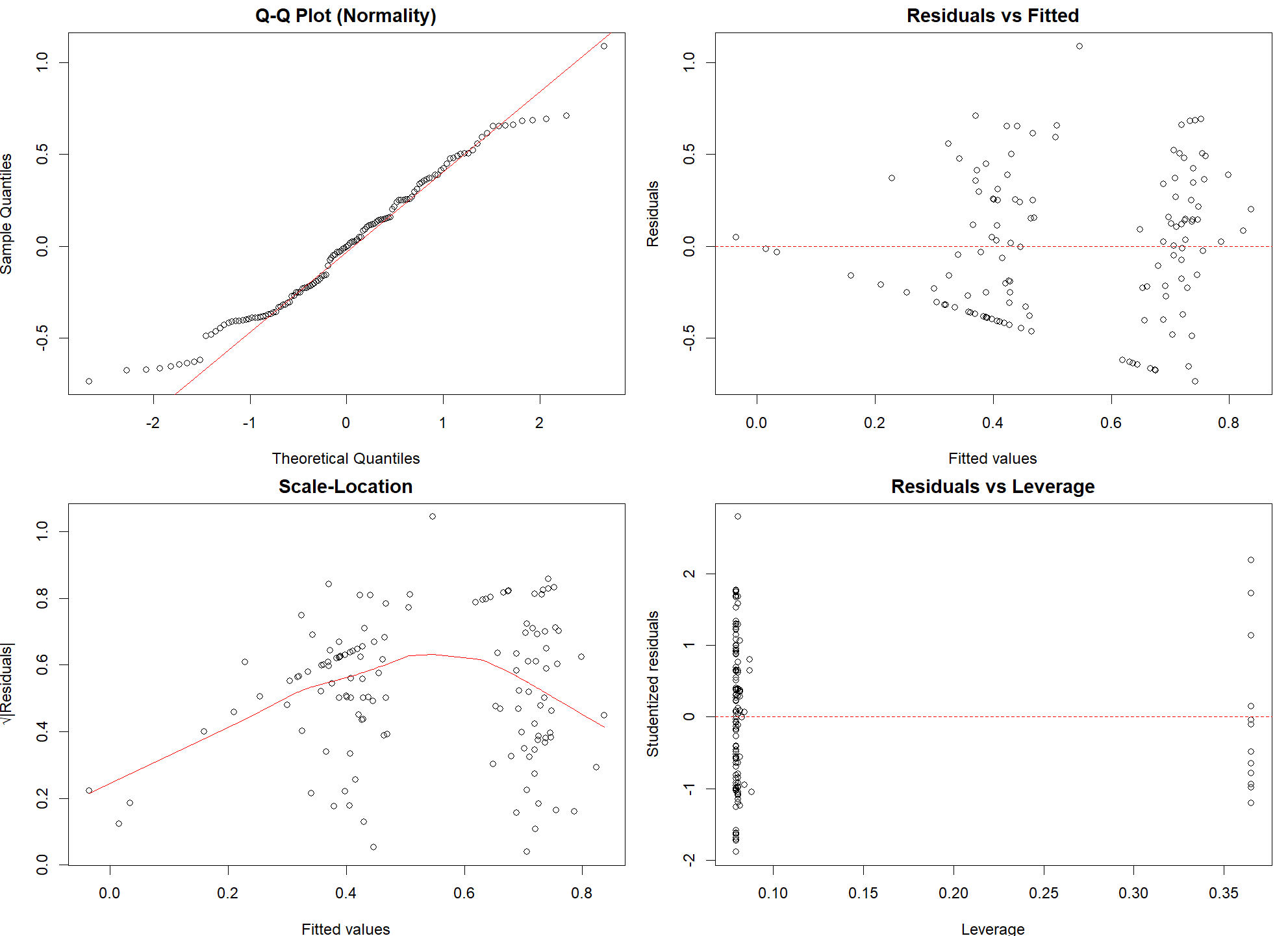

**Figure S3. Diagnostic plots for the linear mixed model (LMM) predicting Shannon diversity.** The Q–Q plot (top left) shows deviations from normality, with long tails in the distribution (Shapiro–Wilk test, p < 0.001). The residuals versus fitted (top right) show a random distribution of points around zero, indicating homoscedasticity. The scale–location plot (bottom left) shows some non-linearity. However, Levene’s test showed homogeneity of variance across groups (p = 0.51). The residuals-versus-leverage plot (bottom right) shows no strong outliers. Together, these diagnostics suggest that although normality assumptions were violated, the homogeneity of variances was satisfied and we therefore deem the use of the LMM as appropriate.

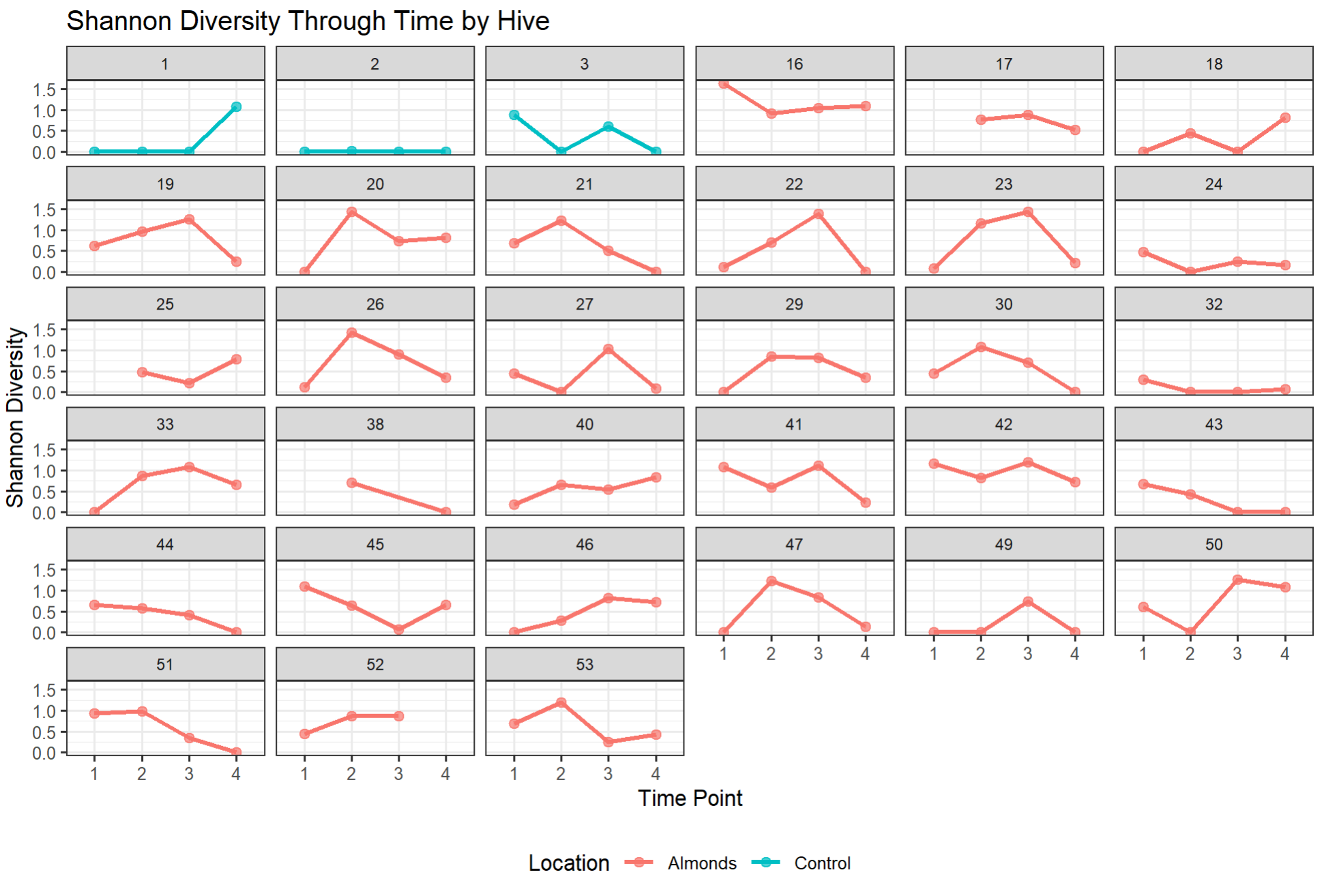

**Fig S4. Shannon diversity index for individual hives plotted through time on line plots.** Blue indicates control hives, whereas pink indicates almond hives. Only hives with viral community data for three out of four time points were plotted.

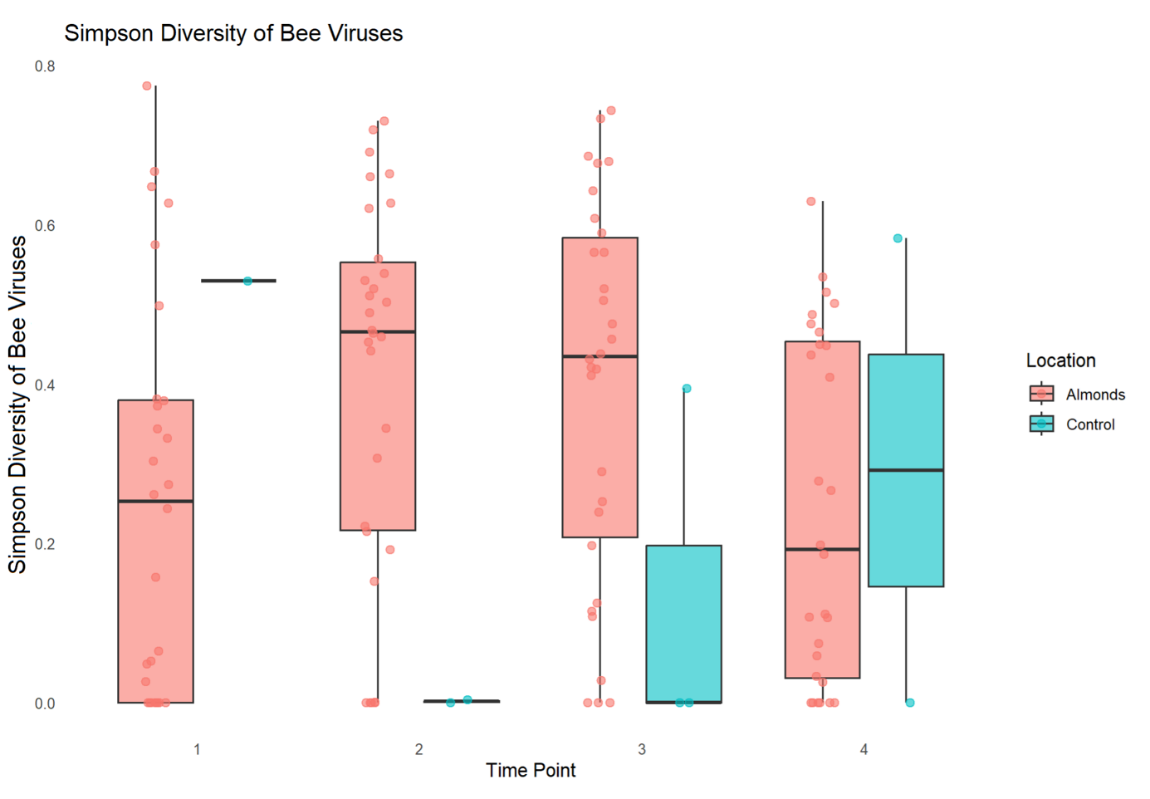

**Figure S5. Simpson diversity of bee viruses across time in almond pollinating and control honeybee colonies.** Boxplots show the normalized Simpson diversity index in almond (pink) and control hives (blue) through time. Points indicate individual hive Simpson diversity values.

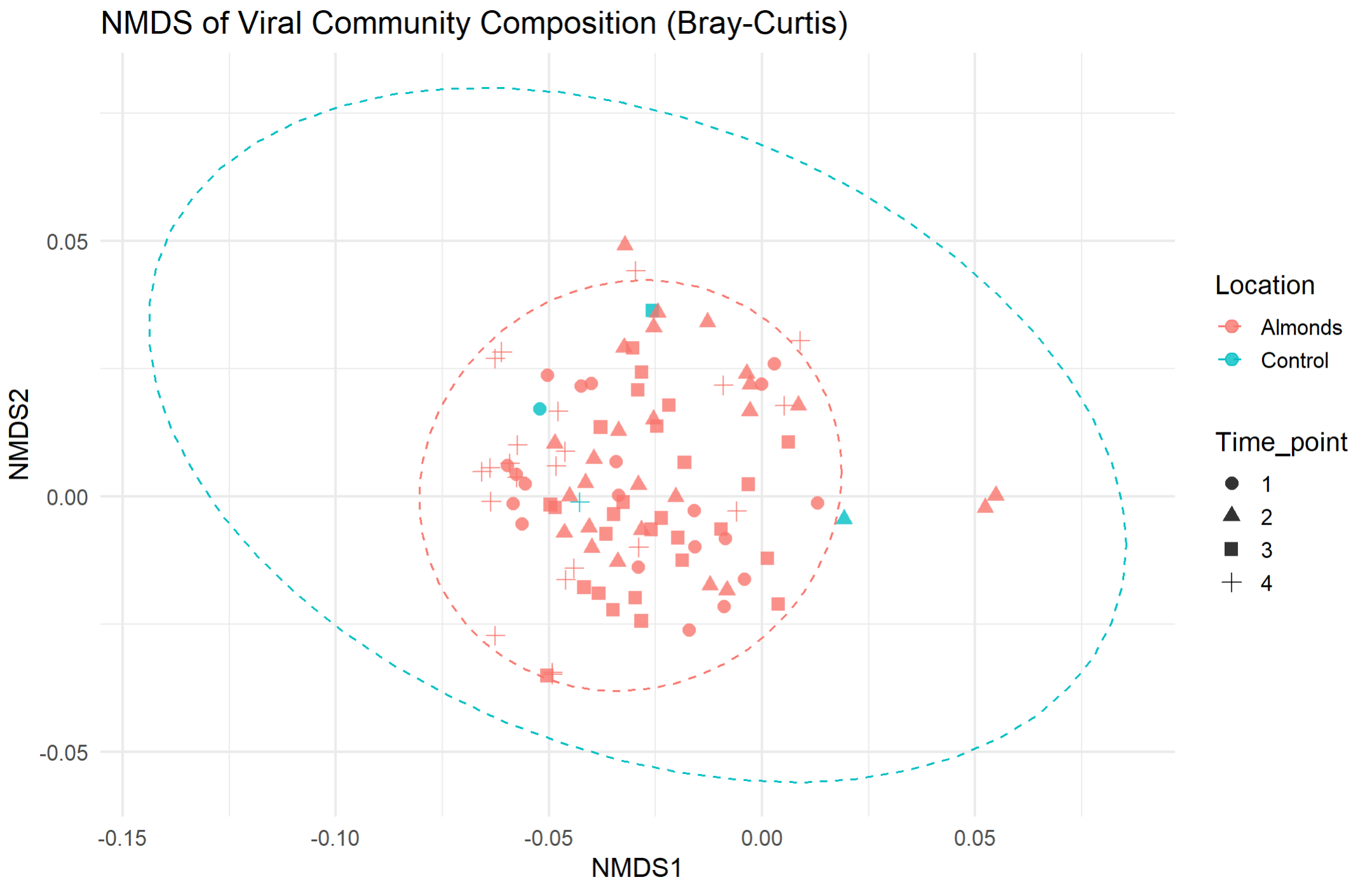

**Figure S6. Non-metric multidimensional scaling (NMDS) of viral community composition across time and location.** Each point represents a hive, colored by the location (almonds versus control), and the shape indicates time point. Viral community dissimilarities were calculated using Bray-Curtis distance on relative abundance data. Ellipses represent 95% confidence intervals around the centroid.

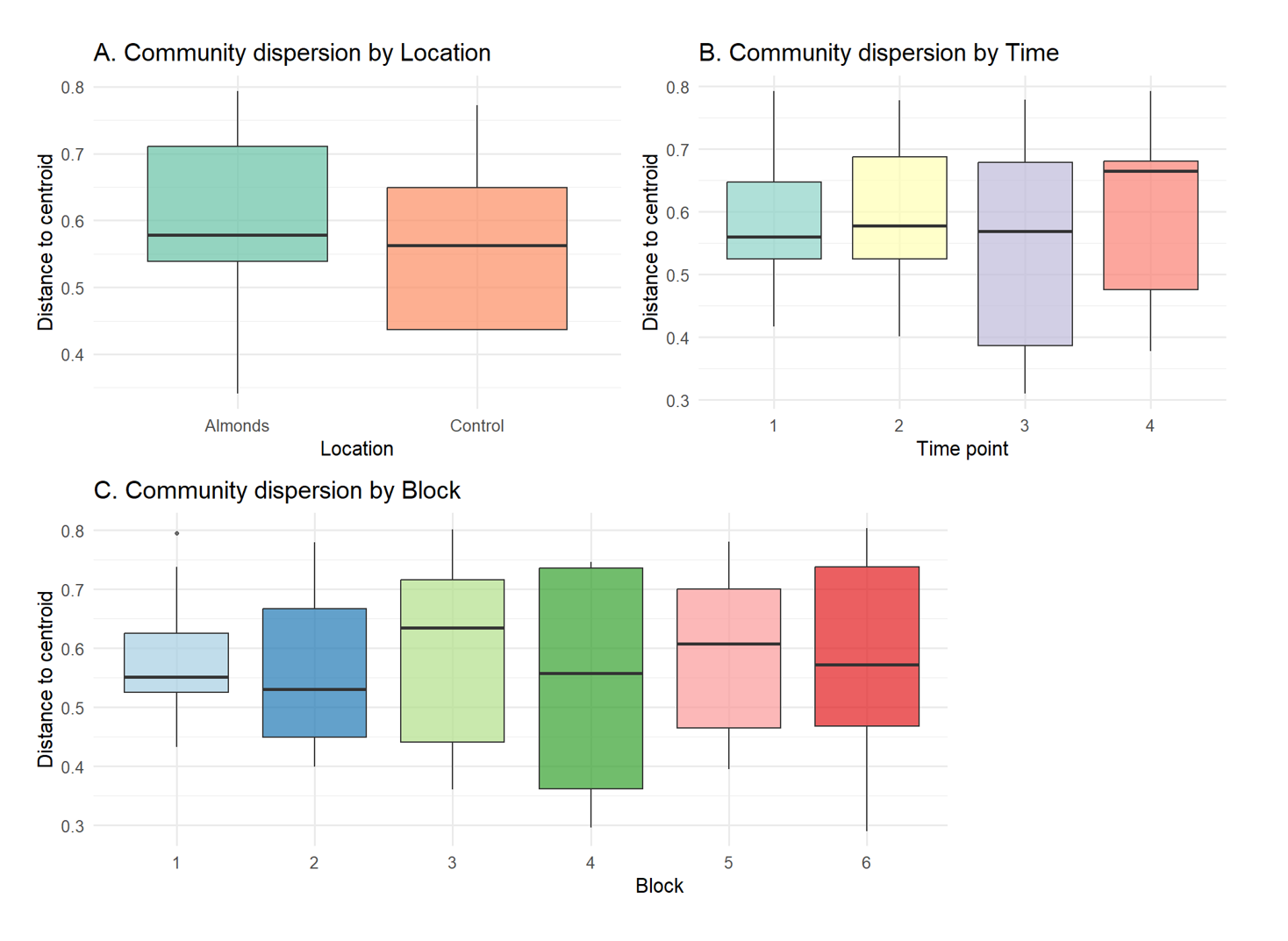

**Fig S7. Multivariate dispersion of viral communities by (A) Location, (B) Time point, and (C) Block, based on Jaccard dissimilarities.** Boxplots show the distance of each sample to its group centroid. PERMDISP tests confirmed no significant differences in dispersion among groups (Location: p = 0.553; Time point: p = 0.248; Block: p = 0.991), indicating comparable within-group variability across factors, and therefore the use of a PERMANOVA model is appropriate.

**Table S1**. Type III ANOVA results for a linear mixed model assessing the effect of time, location, and the interaction between time and location on Shannon diversity with hive number as a random factor.

| **Factor** | **Degrees of freedom** | **Den DF** | **F value** | **P value** |
| --- | --- | --- | --- | --- |
| Time point | 3 | 92.08 | 0.15 | 0.927 |
| Location | 1 | 31.23 | 6.47 | **0.016 *** |
| Time × Location | 3 | 92.08 | 1.66 | 0.18 |

**Table S2.** Post hoc comparisons of Shannon diversity in almond hives in contrast to control hives at each time point with a Benjamini–Hochberg correction for multiple tests.

| **Time point** | **Contrast** | **Estimate** | **SE** | **df** | **t ratio** | **p value** |
| --- | --- | --- | --- | --- | --- | --- |
| 1 | Almonds – Control | 0.136 | 0.254 | 122 | 0.54 | 0.593 |
| 2 | Almonds – Control | 0.704 | 0.253 | 122 | 2.78 | **0.006 **** |
| 3 | Almonds – Control | 0.522 | 0.253 | 122 | 2.06 | **0.042 *** |
| 4 | Almonds – Control | 0.03 | 0.253 | 122 | 0.12 | 0.906 |

**Table S3.** Results of PERMANOVA analyses testing the effects of time point, location, and block on viral community composition using Bray–Curtis and Jaccard dissimilarities.

| **Model + Factors** | **df** | **Sum of Squares** | **R²** | **F** | | **P value** |
| --- | --- | --- | --- | --- | --- | --- |
| **Bray–Curtis (interaction model)** |  |  |  |  | |  |
| Time_point × Location + Block | 4 | 1.571 | 0.03475 | 1.0712 | | 0.3564 |
| Residual | 119 | 43.638 | 0.96525 |  | |  |
| Total | 123 | 45.209 | 1 |  | |  |
| **Jaccard (marginal effects)** |  |  |  |  | |  |
| Time_point | 1 | 0.644 | 0.01754 | 2.2171 | | **0.0293 *** |
| Location | 1 | 0.585 | 0.01595 | 2.0156 | | **0.0503 *** |
| Block | 1 | 0.728 | 0.01983 | 2.5066 | | **0.0180 *** |
| Residual | 120 | 34.848 | 0.94941 |  | |  |
| Total | 123 | 36.705 | 1 |  |  | |
